# Different combinations of insect Na,K-ATPase α- and β-subunit paralogs enable fine tuning of toxin resistance and enzyme kinetics

**DOI:** 10.1101/2020.08.28.272054

**Authors:** Marlena Herbertz, Safaa Dalla, Vera Wagschal, Rohin Turjalei, Marlies Heiser, Susanne Dobler

## Abstract

**Background:** Cardiac glycosides are known to fatally inhibit the Na,K-ATPase throughout the animal kingdom. Several animals, however, evolved target-site insensitivity by substitutions in the otherwise highly conserved cardiac glycoside binding pocket located on the Na,K-ATPase α-subunit. The minimal functional enzyme consist of an α- and a β-subunit, the latter considered mainly as a chaperone responsible for correct folding and membrane integration. We here analyze resistance to cardiac glycosides and kinetic properties of different Na,K-ATPase α/β-combinations of the large milkweed bug, *Oncopeltus fasciatus*. These insects have adapted to high concentrations of cardiac glycosides in their food plants via several rounds of Na,K-ATPase gene duplications followed by differential resistance conferring substitutions and subfunctionalization of the enzymes.

**Results:** To investigate their characteristics we expressed nine combinations of *O. fasciatus* Na,K-ATPase α/β-sunbunits (three each) in *Sf*9 cells and tested them with two structurally distinct cardiac glycosides, calotropin, a host plant compound, and ouabain, a commonly used toxin. Differences in the number and identity of amino acid substitutions in the cardiac glycoside binding site resulted in large differences in activity and toxin resistance of the three α-subunits. The enzymes’ kinetics were also influenced by the β-subunits leading to increased activities (αCβ3) or altered resistances. The host plant toxin calotropin proved to be a much more potent inhibitor than ouabain for the phylogenetically oldest αC based enzymes. This effect was compensated for in the αB and αA based enzymes with αAβ1 having higher resistance against calotropin than against ouabain.

**Conclusion:** The originally higher inhibitory potency of the host compound calotropin supports a coevolutionary escalation of plant defenses and herbivore tolerance mechanisms. For the bugs the possession of multiple paralogs improved adaptation to plant toxins in a stepwise manner and mitigates pleiotropic effects by a compromise between ion pumping activity and resistance.

## Background

The transmembrane Na,K-ATPase, is ubiquitously expressed in animal cells and is essential for maintaining membrane potentials, controlling cell volume and the overall functionality of cells (1). The functional enzyme consists of two subunits, the catalytic α-subunit and an associated β-subunit serving as chaperone for correct folding and membrane integration (2). In vertebrates several paralogs are present for both genes and the enzyme’s characteristics differ depending on the specific combination of α and β-subunits (3,4). An additional modulatory and tissue specific γ-subunit is present in vertebrate tissues (5), but has not been detected in invertebrates. Much less is known about the importance of different α/β-subunit combinations in invertebrates (6). Only in *Drosophila* the tissue distribution and functional roles of some of these combinations have been explored to some extent (7–9). We here explore the functional role of different Na,K-ATPase complexes in the large milkweed bug, *Oncopeltus fasciatus*. In this species, the Na,K-ATPase α1 subunit has undergone three rounds of duplications (10,11) and displays high insensitivity against cardiac glycosides (12,13), which are universal inhibitors of the Na,K-ATPase (1).

Inhibition of the Na,K-ATPase by cardiac glycosides, such as the cardenolides ouabain and calotropin (Fig. 1), depends on a specific binding site located on the extracellular side of the catalytic α-subunit. Upon binding of cardiac glycosides the enzyme is arrested in the phosphorylated state and ion pumping is disrupted (1). Inhibition of the Na,K-ATPase is harmful and can be lethal to almost all animals. There are, however, a few notable exceptions to this: various mammals, frogs, and reptiles prey unharmed on cardiac glycoside-containing toads and a number of phytophagous insects feed - often exclusively - on cardiac glycoside-containing host plants (reviewed in 14,15). Most of these species avoid the toxic effect of cardiac glycosides and exhibit target site insensitivity, by drastically lowering binding affinity of cardiac glycosides to their Na,K-ATPase α-subunit (16,17). Strikingly, identical amino acid substitutions at homologous positions convergently evolved in various, distantly related insect groups and even resemble the substitutions observed in cardiac glycoside-resistant vertebrates (11,17–19).

**Figure 1:**
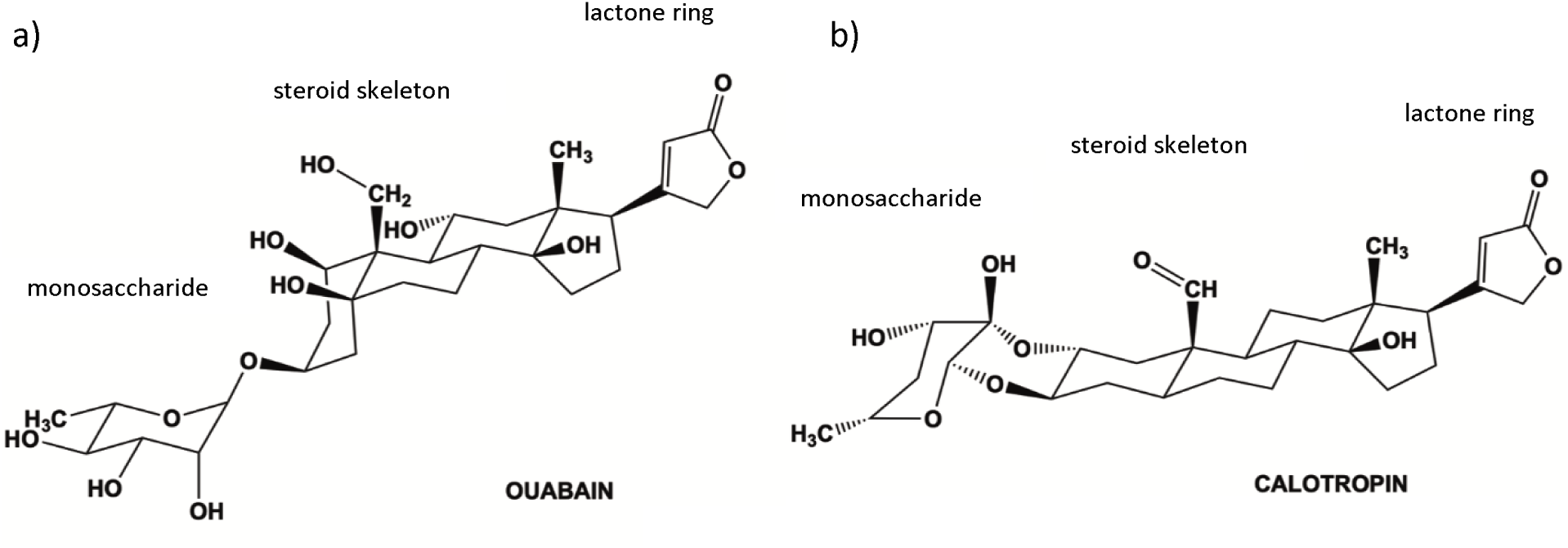
Structure of the cardenolides ouabain (a) and calotropin (b). Both are characterized by a monosaccharide attached to a steroid skeleton at C3 (4 steroid rings: A-D) and a five-membered unsaturated lactone ring at C-17. a) In ouabain the monosaccharide is attached by a single glycosidic link to the steroid rings that are *cis-trans-cis* fused. b) Calotropin is characterized by a doubly linked monosaccharide as well as a *trans-trans-cis* conformation of the steroid rings (figure generated by Daniel Tal, The Weizmann Institute of Science, 2020).

While insect species usually express a single gene copy of the Na,K-ATPase α-subunit (ATPα1) throughout their body, this gene underwent one to three rounds of gene duplications in several cardiac glycoside-exposed species (10,11,16,20). These duplicated ATPα1 copies differ in the number and identity of presumably resistance-causing amino acid substitutions and show tissue specific expression patterns (10,11,20,21).

The large milkweed bug *Oncopeltus fasciatus* (Hemiptera, Lygaeidae) possesses four ATPα1 paralogs (A, B, C, D) that differ in the number and identity of amino acid substitutions (10,11). We previously introduced substitutions observed in these paralogs at four usually highly conserved amino acid sites (Q111, N122, F786, and T797) known to be important for high affinity to cardiac glycosides (22–25) into the sensitive ATPα1 gene of *Drosophila melanogaster* (26,27). The N122H (as observed in ATPα1D of *O. fasciatus*), N122H+T797S (as observed in ATPα1C), Q111T+N122H+F786N (as in ATPα1B) or Q111T+N122H+F78_6_N+T797A (as in ATPα1A) substitutions yielded proteins with stepwise increasing resistance to the cardenolide ouabain. At the same time, enzyme activity decreased with increasing numbers of substitutions (26).

Additional untested substitutions are present in the *O. fasciatus* ATPα1 paralogs and these might influence the characteristics of the enzymes, potentially further increasing resistance but possibly also serving as compensatory mutations mitigating deleterious intra-molecular epistatic effects of previously characterized mutations (28–30). In addition, it has been shown that the β-subunit of the Na,K-ATPase not only acts as a chaperone, guiding the heterodimeric enzyme to the cell membrane, but also influences the behavior of the Na,K-ATPase including its sensitivity to ouabain (2,4,27).

Although ouabain is a commonly used cardiac glycoside in studies of the Na,K-ATPase, it does not occur in host plants of *O. fasciatus*. The *Asclepias* species (Apocynaceae) that the bugs feed on typically contain different types of cardenolides characterized by a *trans-trans-cis* conformation of the steroidal skeleton and a single sugar moiety attached by a cyclic bridge as in the abundant calotropin (15) (Fig. 1). The most biologically relevant test of the bugs’ resistance against cardiac glycosides should thus involve a plant-derived compound like calotropin. To address these issues, we identified and heterologously expressed nine possible *O. fasciatus* Na,K-ATPase α/β combinations and characterized their activities and sensitivities to ouabain and calotropin.

Since activity and resistance of the Na,K-ATPase is determined by the number and identity of amino acid substitutions in the cardenolide binding site located in the α-subunit, (1) we hypothesized that α-subunit paralogs – associated with different β-subunits – differ significantly in their activities and insensitivities to the two cardenolides we tested. (2) We further hypothesized that the different β-subunits modulate the Na,K-ATPase α-subunit paralog behavior influencing their response to the cardenolides. Additionally, although ouabain and calotropin both seem to be strong inhibitors of the Na,K-ATPase, (3) we expected the two cardenolides to differ significantly in their inhibitory potency against the Na,K-ATPase of *O. fasciatus* due to differences in structure and steric conformation. We hypothesized that coevolutionary interaction should lead to higher resistance against the host plant compound calotropin.

## Results

### Bioinformatic analysis

Four potential β-subunits were identified from the *O. fasciatus* transcriptome and according to their similarity to published sequences named β1, β2, β3, and βx (Genbank AccNo xxx-xxx). While β1-3 conform to the standard length of β-subunits of around 900 bp and are recovered in Blast searches as homologues of similarly numbered β-subunits of other insects, βx is much shorter and missing most of the conserved N-terminal and half of the transmembrane region and does not have obvious homologues in other insects (Fig. S1A). This β-subunit was therefore not further considered here. The 5’ end of α1B and α1C could successfully be recovered by RACE protocols (Genbank AccNo xxx, xxx). The only minimally expressed α1D (10) was not further investigated here.

### *Expression of α/β-subunit constructs in* Sf*9 cells*

All nine α/β-subunit constructs (αAβ1, αAβ2, αAβ3, αBβ1, αBβ2, αAβ3, αCβ1, αCβ2, and αCβ3) were successfully expressed in *Sf*9 cells and the corresponding proteins harvested. The western blot shows that similar concentrations were loaded onto the gel (12G10 anti-α-tubulin as positive control), all α-subunits were expressed at similar levels and the deglycosylated β-subunits can be easily discriminated by their specific patterns (Fig. 2).

**Figure 2:**
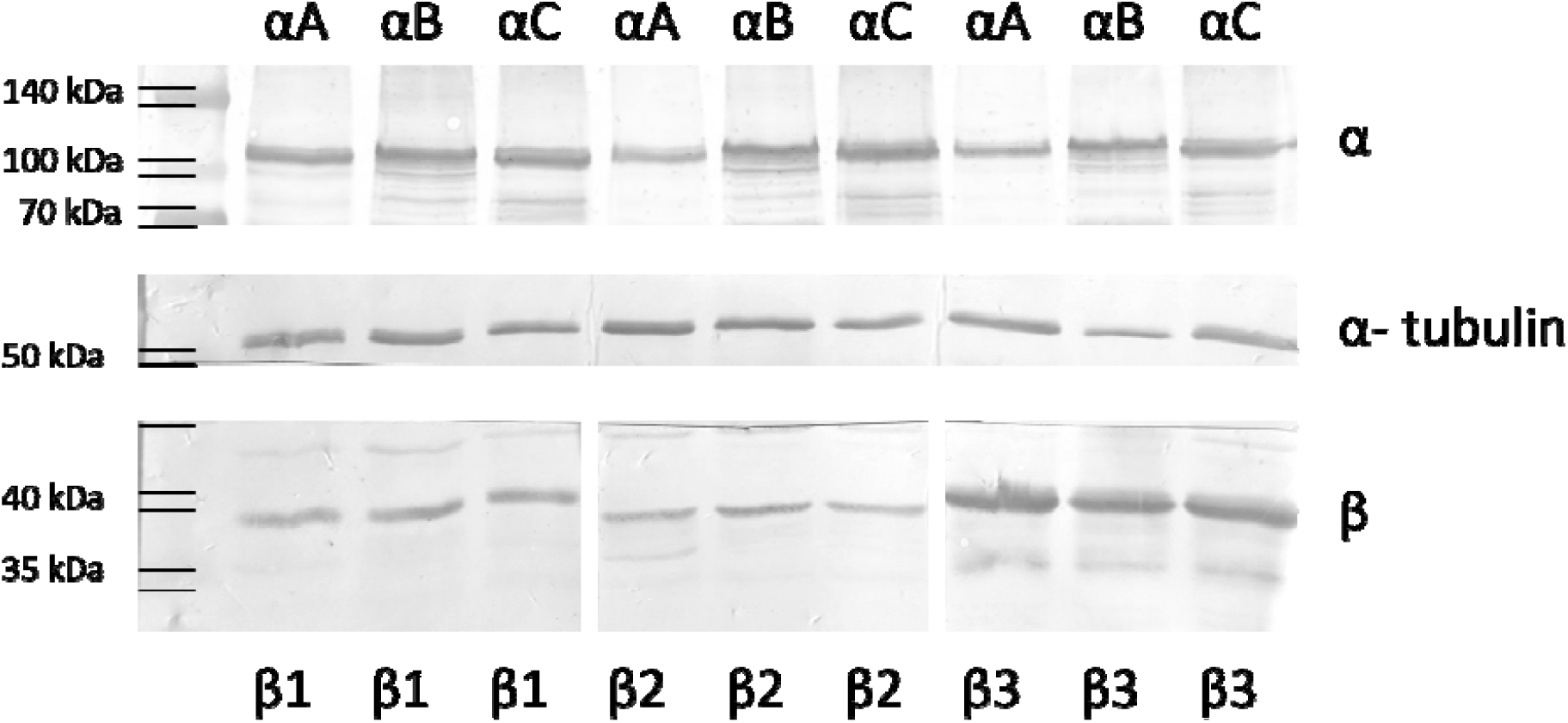
Western blot of nine α/β-complexes of *O. fasciatus* expressed heterologously in cell culture. 30 µg of protein were separated by SDS-PAGE and blotted on a nitrocellulose membrane. The upper membrane slice shows at a size of 110 kDa αA, αB and αC in alternating order, the proteins were stained with the primary antibody α5 (DSHB hybridoma bank). At 55 kDa α-tubulin is shown as a positive control (middle membrane), the protein is stained with the primary antibody 12G10 anti-α-tubulin (DSHB hybridoma bank). Primary antibodies of both proteins were detected with the secondary HRP-conjugated antibody goat-anti mouse (Dianova). The associated β-subunits 1, 2, and 3 were deglycosylated in advance. The lower membrane is split into three slices, the first one shows β1, the second β2, and the third β3. The subunits were stained with β-subunit specific customized primary antibodies (produced by Davids Biotechnology) and HRP-conjugated secondary antibodies goat-anti rabbit (for β2, Dianonva) and goat-anti chicken (for β1 and 3, Dianova). The different β-subunits show bands at weights ranging between 35 and 40 kDa. The bands were visualized with 4-CN. Please note: Signal strength is not comparable between the different β-subunits, due to different binding affinities of the antibodies. Due to the grey scale some bands of the multicolor protein ladder were marked by lines.

### Na,K-ATPase activity of different α/β-complexes

The activities of different α/β-subunit combinations of the Na,K-ATPase were tested *in vitro* to determine whether the amino acid substitutions in the cardenolide binding site or the associated β-subunit influence the catalytic activity of the enzyme. All enzymes showed catalytic activity, but the activity of the α/β-complexes differed significantly from one another (Fig. 3). The α-subunits A and B with their associated β-subunits (1-3) showed significantly lower activity than αC combined with β1, 2 or 3. The activities of αA and αB did not differ significantly from each other regardless of the β-subunit, except for αAβ2. This combination was significantly less active than αB associated with the three β-subunits (Fig. 3). By far the highest activity was measured for αC in association with β3 that had an average turnover of 32.7 nmol Pi per mg Protein per minute and differed significantly from all other enzymes.

**Figure 3:**
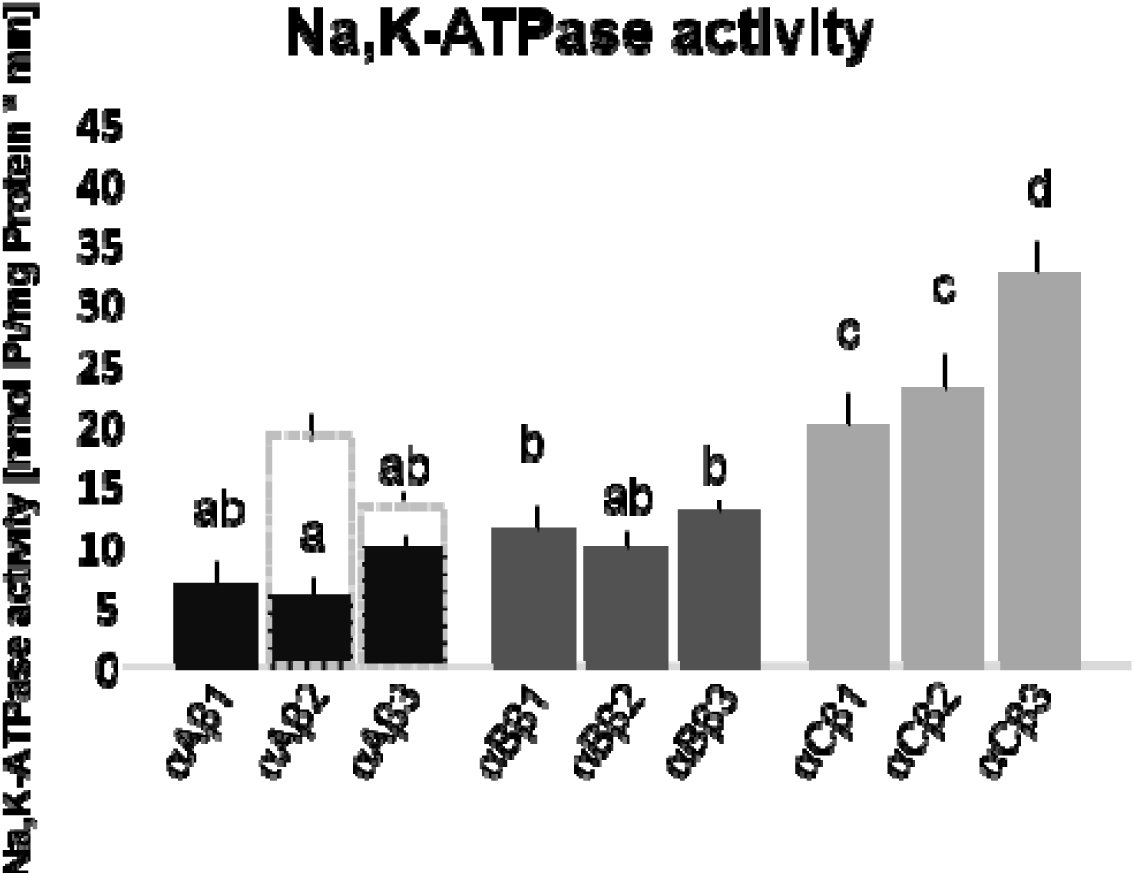
Na,K-ATPase activity [nmol Pi/mg protein*min] of nine α/β-complexes of *O. fasciatus*. Each bar depicts the mean and standard deviation of three biological replicates. Different letters above the bars indicate significant differences between enzymes (ANOVA, F_*8,18*_=77.97, p<0.001). Black bars: αA with three β-subunits and overall activities ranging from **5.5 to 9.3 nmol Pi/mg protein*min**. Bars with a grey dotted outline: data for the αA mimic (THNA) combined with Nrv2.2 and Nrv3 of a previous study (Dalla et al., 2017). Dark gray bars: αB/β-complexes ranging from **10.3 to 13 nmol Pi/mg protein*min**. Light gray bars: αC/β-complexes with activity levels ranging from **19.7 to 32.7 nmol Pi/mg protein*min**. The activity levels of the three αC/β-enzymes are significantly higher than those of the other enzymes, with the highest activity reached by αC combined with β3.

### Influence of α/β-subunit combination on ouabain sensitivity

*In vitro* tests with the cardenolide ouabain revealed large differences in sensitivity of the Na,K-ATPases depending on the number of amino acid substitutions in the cardenolide binding site and was modulated by the identity of the β-subunit (Fig. 4). Yet, when comparing the IC_50_ values only the one of αC associated with β1 was significantly lower than those for the other two αC complexes (ANOVA, F_*2,8*_=8.035, p=0.02, Tukey’s HSD, p=0.035 and p=0.027, for αC β1 vs β2 and β1 vs β3, respectively; Fig. 4c, Table 1) whereas the IC_50_ values of the three αA or αB complexes did not differ significantly.

**Figure 4:**
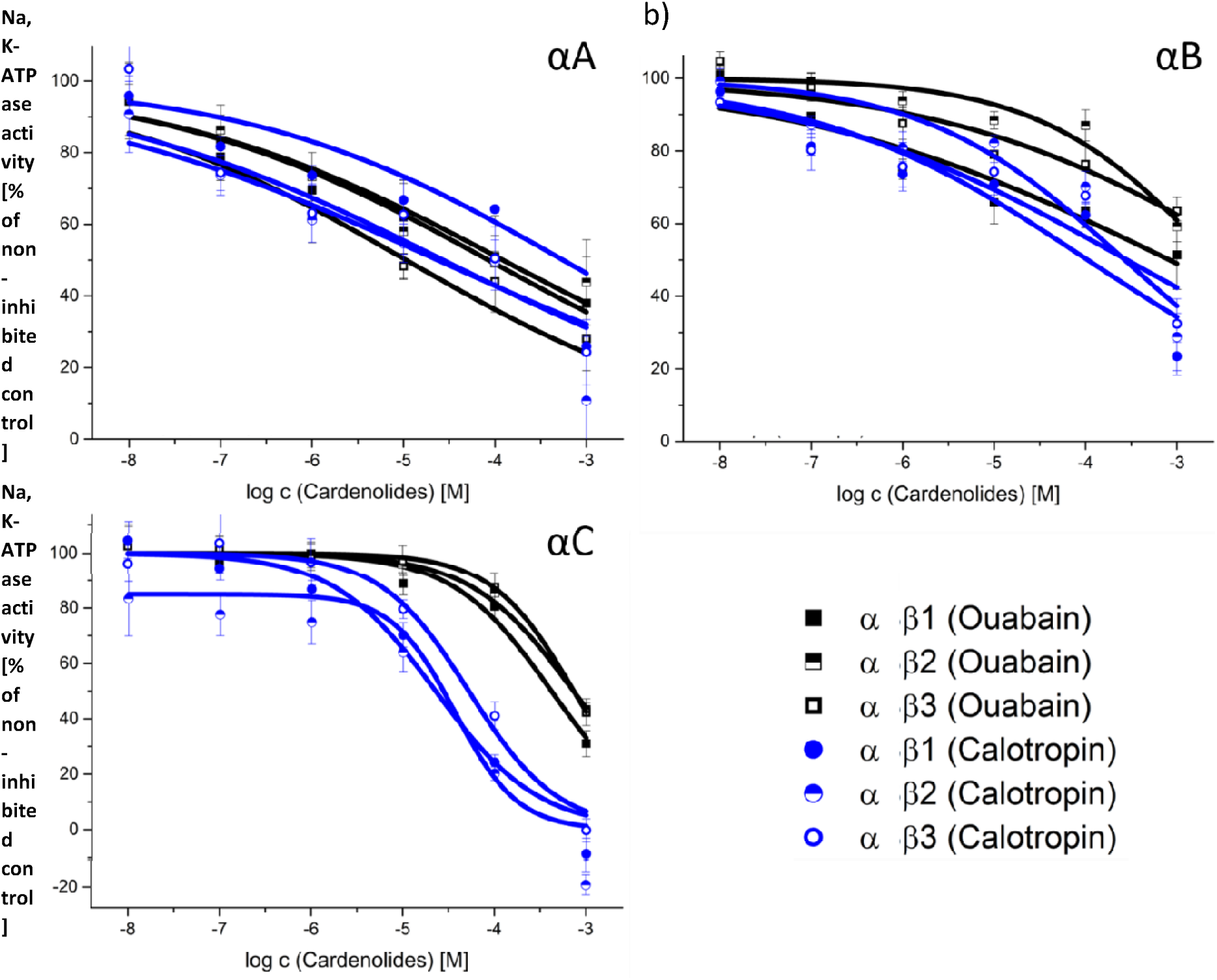
***In vitro* inhibition by calotropin and ouabain of nine α/β-complexes of *O. fasciatus* expressed heterologously in cell culture. Dose response relationships between increasing cardenolide concentrations and residual enzyme activities were expressed in % of the non-inhibited control (calculated with Origin Pro 9.1G).** Each data point represents the mean of three biological replicates and standard deviation. Effect of calotropin (blue curves and round symbols) and ouabain (black curves and square symbols) a) on αA, b) on αB, and c) on αC combined each with β1, 2, and 3. Please note the different scales.

**Table 1:** Mean IC_50_ values and Hill slopes of different Na,K-ATPase α/β-complexes (3 biological replicates with standard deviation (± s.d.)) obtained from dose response curves upon inhibition with ouabain and calotropin (Origin-Pro 9.1G).

| $\alpha/\beta$-<br>combination | $IC_{50}$ values [M]<br>Ouabain (± s.d.) | Hill slope<br>(± s.d.) | $IC_{50}$ values [M]<br>Calotropin<br>(± s.d.) | Hill slope<br>(± s.d.) |
| --- | --- | --- | --- | --- |
| $\alpha A\beta 1$ | 4.148E-04<br>(± 6.541E-04) | -0.221<br>(± 0.047) | 1.211E-04<br>(± 4.443E-05) | -0.271<br>(± 0.03) |
| $\alpha A\beta 2$ | 3.471E-04<br>(± 3.598E-04) | -0.234<br>(± 0.013) | 1.449E-05<br>(± 5.994E-06) | -0.269<br>(± 0.076) |
| $\alpha A\beta 3$ | 2.607E-05<br>(± 2.767E-05) | -0.260<br>(± 0.044) | 6.959E-05<br>(± 7.54E-05) | -0.27<br>(± 0.036) |
| $\alpha B\beta 1$ | 2.089E-03<br>(± 2.515E-03) | -0.213<br>(± 0.039) | 1.108E-04<br>(± 3.423E-05) | -0.293<br>(± 0.044) |
| $\alpha B\beta 2$ | 2.777E-03<br>(± 9.676E-04) | -0.462<br>(± 0.055) | 3.281E-04<br>(± 2.119E-04) | -0.398<br>(± 0.066) |
| $\alpha\text{B}\beta 3$ | 7.09E-03<br>($\pm 3.119\text{E-}03$) | -0.261<br>($\pm 0.045$) | 2.944E-04<br>($\pm 8.195\text{E-}05$) | -0.254<br>($\pm 0.071$) |
| $\alpha\text{C}\beta 1$ | 4.274E-04<br>($\pm 8.104\text{E-}05$) | -0.869<br>($\pm 0.071$) | 2.402E-05<br>($\pm 4.636\text{E-}06$) | -0.88<br>($\pm 0.029$) |
| $\alpha\text{C}\beta 2$ | 7.369E-04<br>($\pm 1.001\text{E-}04$) | -0.869<br>($\pm 0.093$) | 3.699E-05<br>($\pm 5.517\text{E-}06$) | -1.289<br>($\pm 0.198$) |
| $\alpha\text{C}\beta 3$ | 7.579E-04<br>($\pm 1.476\text{E-}04$) | -0.988<br>($\pm 0.35$) | 5.594E-05<br>($\pm 8.55\text{E-}06$) | -0.912<br>($\pm 0.098$) |

Although analyzing differences in IC_50_ is a standard approach, the IC_50_ values only reflect one point of the dose response curve. We therefore aimed to get a better insight into the overall responses of the Na,K-ATPases to increasing ouabain concentrations by analyzing the differences between the full dose response curves of enzymes with the same α-subunit as well as enzymes with the same β-subunit (Table 2).

**Table 2:** Comparison between the dose response curves of different α/β-complexes inhibited by ouabain and calotropin analyzed with Origin-Pro 9.1G. Significant differences are highlighted in bold.

| Data comparison | F-statistic | F-statistic |
| --- | --- | --- |
| $\alpha/\beta$-combinations | Ouabain | Calotropin |
| $\alpha\text{A}\beta 1 - \alpha\text{A}\beta 2$ | $F_{2,8} = 0.074, p = 0.929$ | $F_{2,8} = 1.319, p = 0.320$ |
| $\alpha\text{A}\beta 1 - \alpha\text{A}\beta 3$ | **$F_{2,8} = 5.218, p = 0.035$** | $F_{2,8} = 0.728, p = 0.512$ |
| $\alpha\text{A}\beta 2 - \alpha\text{A}\beta 3$ | **$F_{2,8} = 6.092, p = 0.025$** | $F_{2,8} = 0.079, p = 0.925$ |
| $\alpha\text{B}\beta 1 - \alpha\text{B}\beta 2$ | **$F_{2,8} = 10.981, p = 0.005$** | $F_{2,8} = 2.607, p = 0.134$ |
| $\alpha\text{B}\beta 1 - \alpha\text{B}\beta 3$ | $F_{2,8} = 3.781, p = 0.070$ | $F_{2,8} = 0.724, p = 0.514$ |
| $\alpha\text{B}\beta 2 - \alpha\text{B}\beta 3$ | $F_{2,8} = 0.455, p = 0.650$ | $F_{2,8} = 0.655, p = 0.545$ |
| $\alpha\text{C}\beta 1 - \alpha\text{C}\beta 2$ | $F_{2,8} = 0.745, p = 0.505$ | $F_{2,8} = 1.887, p = 0.213$ |
| $\alpha C\beta 1 - \alpha C\beta 3$ | $F_{2,8} = 1.535, p = 0.273$ | $F_{2,8} = 3.402, p = 0.085$ |
| $\alpha C\beta 2 - \alpha C\beta 3$ | $F_{2,8} = 1.080, p = 0.384$ | $F_{2,8} = 2.217, p = 0.171$ |
| $\alpha A\beta 1 - \alpha B\beta 1$ | $F_{2,8} = 1.039, p = 0.397$ | $F_{2,8} = 0.554, p = 0.595$ |
| $\alpha A\beta 1 - \alpha C\beta 1$ | **$F_{2,8} = 22.729, p < 0.001$** | **$F_{2,8} = 8.724, p = 0.010$** |
| $\alpha B\beta 1 - \alpha C\beta 1$ | **$F_{2,8} = 11.975, p = 0.004$** | **$F_{2,8} = 8.795, p = 0.010$** |
| $\alpha A\beta 2 - \alpha B\beta 2$ | **$F_{2,8} = 57.311, p < 0.001$** | **$F_{2,8} = 9.763, p = 0.007$** |
| $\alpha A\beta 2 - \alpha C\beta 2$ | **$F_{2,8} = 4.835, p = 0.042$** | **$F_{2,8} = 6.422, p = 0.022$** |
| $\alpha B\beta 2 - \alpha C\beta 2$ | $F_{2,8} = 2.353, p = 0.157$ | **$F_{2,8} = 13.589, p = 0.003$** |
| $\alpha A\beta 3 - \alpha B\beta 3$ | **$F_{2,8} = 32.590, p < 0.001$** | **$F_{2,8} = 37.553, p < 0.001$** |
| $\alpha A\beta 3 - \alpha C\beta 3$ | **$F_{2,8} = 33.295, p < 0.001$** | **$F_{2,8} = 19.271, p < 0.001$** |
| $\alpha B\beta 3 - \alpha C\beta 3$ | $F_{2,8} = 1.614, p = 0.258$ | **$F_{2,8} = 8.239, p = 0.011$** |

We found that α-subunits associated with different β-subunits behaved differently when subjected to ouabain. The enzyme αAβ3 was significantly more sensitive to the cardenolide than the other two αA complexes (Table 2, Fig. 4a), and αBβ1 was inhibited more strongly by ouabain than αBβ2 (Table 2, Fig. 4b). Nevertheless, the reaction of enzymes with identical α-subunits was more uniform than the one of Na,K-ATPases with identical β but different α-subunits. Most of the latter enzymes differed significantly in their reactions, reflecting their varying insensitivity levels against ouabain (Fig. 4 a-c, Table 2).

### Influence of α/β-subunit association on calotropin sensitivity

Testing the inhibitory effect of the host plant cardenolide calotropin on the recombinant Na,K-ATPases resulted in total inhibition of the most calotropin-sensitive αC/β-combinations at the highest calotropin concentration. Furthermore, the three αB based enzymes turned out to be most resistant against calotropin. As for the inhibition by ouabain only the IC_50_ values of αC associated with the three different β-subunits differed significantly from each other, αC associated with β3 was significantly more insensitive to calotropin than αC associated with β1 and β2 (ANOVA, F_*2,8*_=18.554, p=0.003, Tukey’s HSD, p=0.002 and p=0.027, for αCβ3 versus αCβ1 and αCβ3 versus αCβ2, respectively; Fig. 4c).

When comparing the full dose response curves of the constructs (Table 2) neither the three αA combinations differed significantly from one another nor the αB or αC based enzymes, indicating a similar response to calotropin exposition independent of the associated β-subunit (Fig. 4 a-c). In contrast, the inhibition by calotropin of the three α-subunits combined with the same β-subunits was significantly different (from high to low inhibition: αC> αA> αB; Fig. 4a-c). The curves of αAβ1 and αBβ1 did not differ significantly from one another, since αAβ1 tends to be the most resistant combination of the three αA enzymes and αBβ1 the most sensitive of the three αB enzymes (Fig. 4a & b).

### Comparison of both cardenolides

Calotropin inhibited especially the three αC enzymes to a much higher extent than ouabain as judged by a comparison of IC_50_ values (ANOVA, F_*5,17*_=57.25882, p<0.001, Tukey’s HSD, p<0.001 for all αC based enzymes; Fig. 4c, Table 1). αCβ1 was inhibited 18-fold, αCβ2 20-fold, and αCβ3 14-fold more strongly by calotropin than by ouabain. When only comparing IC_50_ values αB associated with β3 was also significantly more strongly inhibited by calotropin than ouabain, resulting in an approximately 24-fold higher IC_50_ (ANOVA, F_*5,17*_=7.54487, p=0.00205, Tukey’s HSD, p=0.00357; Fig. 4b, Table 1) while the IC_50_ values of the two cardenolides did not differ significantly for αBβ1 and αBβ2 (Tukey’s HSD, p=0.70595 and p=0.51146, respectively; Fig. 4b, Table 1). Comparison of the two cardenolides resulted in no significant differences between IC_50_ values of αA associated with any of the three β-subunits (ANOVA, F_*5,16*_=1.85583, p=0.18241; Fig. 4b, Table 1).

However, when comparing the full dose response curves of each enzyme inhibited by either of the cardenolides, we found that αAβ1 was significantly more insensitive against calotropin than against ouabain, reflected by a flat descending curve at the top (Fig. 4a, Table 3). For the αC-based enzymes the stronger inhibition by calotropin remained evident, while for enzymes with αB no difference in inhibitory potency between the two cardenolides was found (Table 3).

**Table 3:**
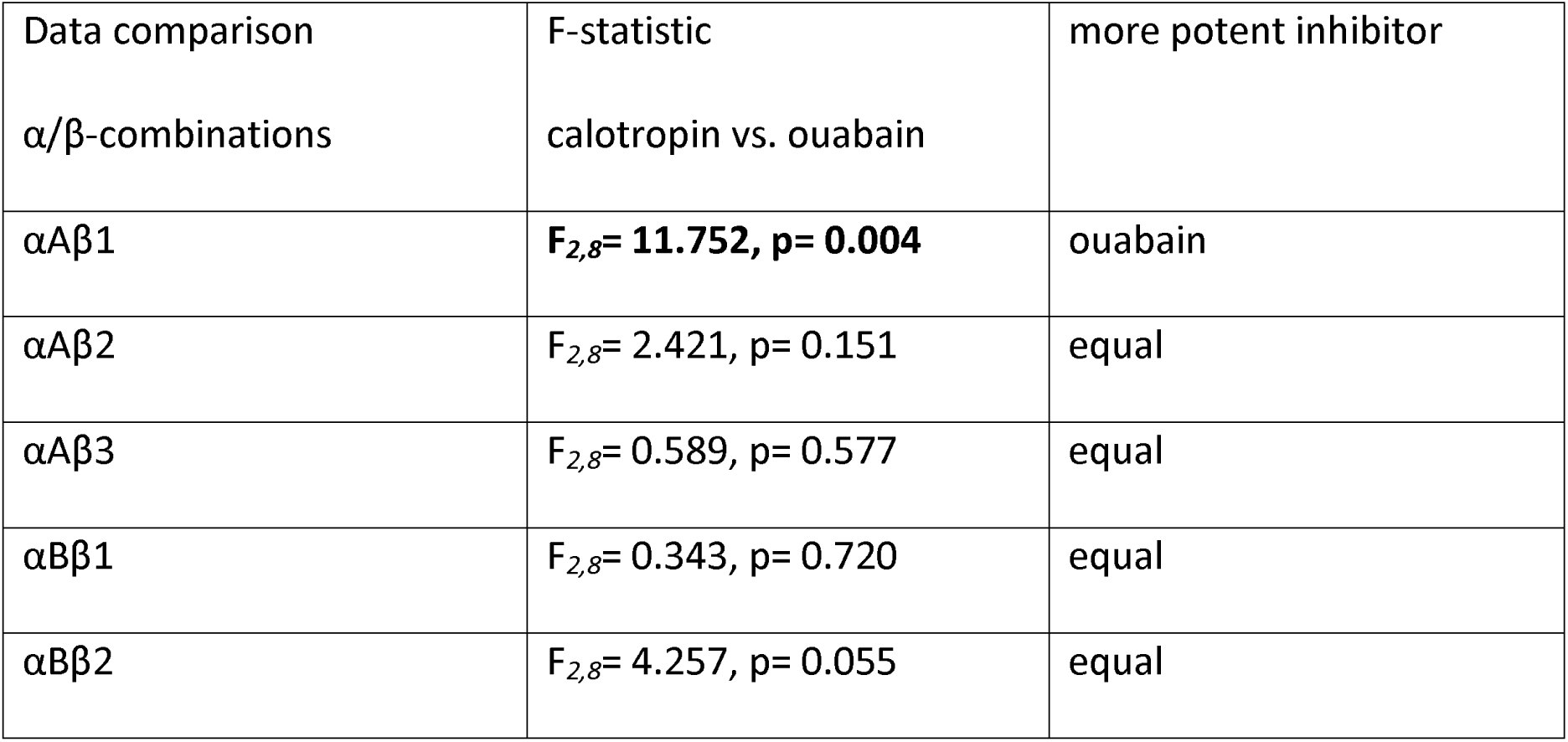
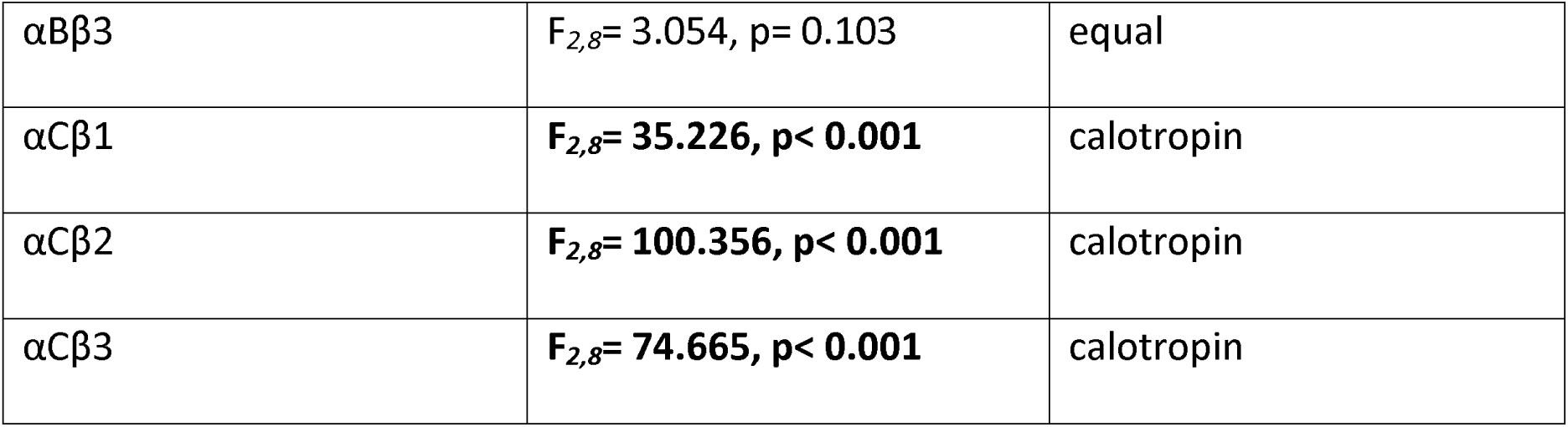
Comparison between dose response curves for individual α/β-complexes inhibited by ouabain versus calotropin analyzed with Origin-Pro 9.1G. Significant differences are highlighted in bold.

## Discussion

Here we have identified the full length coding sequences of three Na,K-ATPase α- and three full length β-subunits of *O. fasciatus* and functionally characterized all nine possible recombinant α/β enzyme complexes. These nine functional enzymes strongly differed in their activity levels. We found that enzyme complexes with αC had a significantly higher activity compared to the other constructs. This perfectly fits with the fact that αC is mainly expressed in the nervous tissue of *O. fasciatus* (11,21) where the Na,K-ATPase plays a crucial role in maintaining electric potentials that guarantee the excitability of neurons. Consequently, a high ionic pumping activity is indispensable. Two of our nine α/β-complexes clearly stand out amongst the other enzyme complexes: αAβ2 and αCβ3. The activity of αAβ2 is reduced in comparison to the other αA and αB constructs that all show similar activity levels. Therefore, we suppose that αAβ2 is not a predominant enzyme *in vivo*. In contrast, the activity of αC combined with β3 is significantly higher and would therefore fit well with a natural occurrence of this complex in the nervous tissue. Data of an ongoing proteomic study support this assumption and show that the subunits αC and β3 are most abundant in the nervous tissue of *O. fasciatus* (M. Herbertz et al., unpublished data).

Regardless of the strikingly high overall resistance to inhibition by cardenolides that Na,K-ATPase preparations of *O. fasciatus* display (12,13,26) we found that the individual α/β-combinations strongly differed in their resistance against the two cardenolides ouabain and calotropin. These two cardenolides differ in their structure, steric conformation and by their ability to permeate membranes, as reflected by their lipophilicity. Ouabain is produced in plants of the genus *Acokanthera* and in seeds of *Strophanthus gratus* (both Apocynaceae), it is commercially available, very polar and routinely used in studies on Na,K-ATPase resistance to cardiac glycosides (1,12,23,25,27). Calotropin, on the other hand, is a common compound in plants of the tribe Asclepiadeae (Apocynaceae) to which the host plants of *O. fasciatus* belong. The two cardenolides differ in the conformation of the steroid rings, which is supposed to determine inhibitory potency. The A/B and C/D rings of ouabain, and many other cardenolides, are *cis*-fused and the B/C ring is *trans*-fused (*cis-trans-cis*) (15,31). In contrast calotropin and other cardenolides of *Asclepias* species, have a *trans*-*trans*-*cis* conformation (15) that is predicted to lower the toxicity (32) and the binding affinity of the cardenolide (31). This may, however, be compensated by the doubly-linked sugar moiety, typical of *Asclepias* cardenolides (33, 34). A recent study testing various cardiac glycosides against the Na,K-ATPase of the monarch butterfly, *Danaus plexippus*, which is also specialized on *Asclepias* species, actually found calotropin and its isomer calactin to be the strongest tested inhibitors, with calotropin inhibiting the monarch’s Na,K-ATPase 60-fold more strongly than ouabain (35).

In contrast to the monarch butterfly, there is evidence that the coevolutionary association between the milkweed bugs and their host plants led to lowered sensitivity towards calotropin. While the phylogenetically older αC-subunit (11, 26) was roughly 20-times more strongly inhibited by calotropin than by ouabain not only the overall sensitivity to both cardenolides but also the relative inhibition by calotropin compared to ouabain drastically decreased in the phylogenetically more derived αA- and αB-subunits. Dose-response curves of αB-based enzymes did not significantly differ with respect to inhibition by either cardenolide whereas the αAβ1 complex was even more resistant to calotropin than to ouabain. This argues in favor of a specific adaptation to the host plant compounds over the course of the evolutionary interaction between the milkweed bugs and their host plants.

The very pronounced insensitivity of the αA- and αB-subunits of *O. fasciatus* to both cardenolides fits well with their abundant expression in the Malpighian tubules (21) where the enzymes are directly exposed to toxins in the hemolymph and a very efficient reabsorption of cardenolides from the primary urine takes place (36). In contrast, the least sensitive but most active αC is the dominant Na,K-ATPase α-subunit in the nervous tissue where it is protected from intruding cardenolides by the perineurium (blood-brain-barrier) which controls xenobiotica access to the nervous system by a tight network of septate junctions and active efflux carriers (37).

A comparison of the present results with our previous studies (26,27) on *O. fasciatus* Na,K-ATPase mimics reveals good concordance regarding our conclusions about enzyme activities. We previously introduced one, two, three or four of the potentially resistance-conferring substitutions into the α*-*subunit of *D. melanogaster* to mimic the known α1-gene copies of *O. fasciatus* (in this order: αD, αC, αB, αA-mimic; c.f. 10,11) and combined them with two *D. melanogaster* β-subunits, Nrv2.2 and Nrv3 (26,27). The overall behavior of these mimics strongly resembled the present data for the original *O. fasciatus* Na,K-ATPase complexes. Most strikingly, the activity levels of the αA and αB mimics were significantly lower than of the Drosophila wildtype enzyme, the αC or the αD (N122H) mimics. The two Drosophila β-subunits also modulated the activity: the association with Nrv3 strongly reduced the activity of all mimics and the Drosophila wildtype enzyme when compared to those bearing Nrv2.2. In the present study, such a large influence of the β-subunit on ion pumping activity was observed only for the αC-subunit while the effect was less pronounced for the αA and αB-subunits.

The β-subunits also strongly modulated the resistance of the enzymes against ouabain, but their effect was specific to the α-subunits they were combined with. A combination with Nrv3 instead of Nrv2.2 previously lead to lowered resistance in the αA- and αB-mimic and increased resistance in the αC-mimic (27). In the present study, resistance changed in the same direction with β1, β2, or β3 for αA (decreasing) and αC (increasing IC_50_) but also lead to increased IC_50_-values for αB.

The most striking differences, however, appear when comparing the overall ouabain-sensitivity of the original enzyme combinations with the previously constructed mimics (27). While the mimics of the *O. fasciatus* αA-copy (featuring the substitutions Q111T,N122H,F786N,T797A) were by far the most insensitive constructs followed by the αB-mimics (with Q111T,N122H,F786N), and the more sensitive αC-mimics (N122H,F786S), the original αA-constructs tested here are at least five times less sensitive than αB-constructs. On the other hand, the original αC-constructs are 10-times more resistant than predicted from the experiments with the αC-mimics.

These differences in sensitivity between mimics and original enzymes strongly point to the importance of additional substitutions in the *O. fasciatus* Na,K-ATPase α1-paralogs. When constructing the mimics, we focused on the four most conspicuous amino acid exchanges in the binding pocket of the Na,K-ATPase that were previously shown functionally to be essential for cardiac glycoside binding (25,38,39). However, there are additional amino acid substitutions in the cardiac glycoside binding pocket of the Na,K-ATPase α1-genes of *O. fasciatus* that may further increase their resistance (11,19). These and further additional amino acid exchanges that at first appear inconsequential may be of importance in modulating the effects of the resistance conferring substitutions or ameliorating their pleiotropic effects (28-30).

## Conclusion

Resistance to specific cardiac glycosides, enzyme activity and the emerging knowledge on tissue distribution of the nine α/β-complexes of the large milkweed bug investigated here further support a scenario of stepwise adaptation to the bugs’ toxic host plants: The duplications leading to three paralogs of the α1-subunit gene that can be combined with three functional β-subunits enabled a functional diversification of the resulting Na,K-ATPases allowing for tissue specific adaptations. The high activity of the αC/β-complexes is suited for efficient ion transport necessary for excitability of nervous tissue but trades off against lower resistance to high concentrations of cardiac glycosides. These phylogenetically older complexes were relatively more susceptible to the host plant toxin calotropin while the younger αA- and αB-subunits were overall more resistant and relatively less sensitive to calotropin and thus suited for expression in toxin exposed tissues like the Malpighian tubules. These physiological details provide new fascinating insights into the continued escalation of defense and counter-defenses between insects and their poisonous host plants.

## Material and methods

### Identification of sequences

Two of the three previously published gene copies of the Na,K-ATPase α1 gene (11) (α1B Acc.No. JQ771519, α1C Acc.No. JQ771518) were incomplete at the 3’ end. Both missing ends could be recovered by RACE PCR (SMARTer RACE kit, Takara Bio, Mountain View, CA, USA).

Previously published transcriptomic data of *O. fasciatus* (11) were used to identify the full-length coding sequences of the Na,K-ATPase β-subunits. The raw data were trimmed and assembled with Trinity (40) as described in Meinzer et al. (41). *O. fasciatus* β-subunit sequences were recovered using published orthologous sequences from *Locusta migratoria* (Orthoptera; Acc.No. KF813098-KF813100) and *Blatta germanica* (Blattodea; Acc.No. HE819405) in tblastn searches against the transcriptome. *De novo* assembly of the raw reads and quality control was performed with the program CLC-Genomics Workbench (Qiagen, Hilden, Germany). The resulting *contigs* were screened for open reading frames (ORFs) with a length of roughly 900 bp (the consensus length of the conserved Na,K-ATPase β-subunit) using the program VectorNTI (Thermo Fisher, Darmstadt, Germany). Homology of the *contigs* were assigned by a further round of tblastx searches against published sequences in genbank.

### Construction of recombinant baculoviruses

The coding sequences of the Na,K-ATPase α- and β-subunits genes (α1A, α1B, α1C, β1, β2 and β3) were amplified from cDNA transcribed with Superscript III (Thermo Fisher) from RNA extracted from nervous tissue, gut, testes or muscles with the RNAmagic-kit (Bio-Budget Technologies GmbH, Krefeld, Germany). For downstream cloning all subunits were amplified with the Pfu-Phusion Polymerase (Thermo Scientific) and primers including the full coding sequence and incorporating suitable restriction sites at the 5’ end.

PCR products were restricted with the chosen enzymes (Thermo Fisher), gel extracted (QIAquick gel extraction kit, Qiagen) and ligated into pFastBac-Dual vectors (Thermo Fisher). The final plasmids had an α-subunit under the control of the polyhedrin promoter (PH) and a β-subunit under the p10 promoter. Recombinant baculoviruses were generated as previously described (27) and transfected into *Sf*9 cells to obtain a first (P1) and second (P2) generation of recombinant virus stock. Uninfected *Sf*9 cells have no or negligible Na,K-ATPase abundances so that contaminating endogenous Na,K-ATPase activities can be excluded (42).

### Production of recombinant proteins

Production of different recombinant Na,K-ATPase α-β-subunit constructs started from infection of Sf9 cells with P2 virus stock and followed our previously published protocol (27). The cells were harvested by centrifugation, the pellet resuspended and sonicated and cell debris removed by centrifugation. Cell membranes were recovered by ultracentrifugation of the supernatant and redissolved in pure water (ROTIPURAN® p.a., ACS water; Roth). The concentrations of the resuspended protein pellets were determined by Bradford assay (Sigma-Aldrich, Taufkirchen, Germany) using bovine serum albumin (BSA) as a standard.

### SDS-PAGE/ Western blotting

Specific binding of customized β-specific anitbodies (Davids Biotechnology, Regensburg, Germany) was additionally validated in our lab by applying the antibodies on two experimental setups. First, the specificity of the antibodies was tested with heterologously expressed *O. fasciatus* Na,K-ATPase protein and second, with four different tissue extracts (muscles, Malpighian tubules, ovaries and nervous tissue). The α1/β1, α1/β2, α1/β3, and α1/βx cell protein construct extracts (4× 30 µg) and muscles, Malpighian tubules, ovaries, and nervous tissue extract (4× 35 µg for each tissue) were run on a SDS-PAGE followed by western blotting as described below. All four Na,K-ATPase constructs and tissues were immunostained with each of the specific β-antibodies. These stainings verified that no unspecific binding occurs (Fig. S2 & S3), but that the β-specific antibodies solely detect their target-subunit in all tested samples.

Thirty µg of the heterologously expressed proteins were deglycosylated with endoglycosidase H following the manufacturer’s protocol (Promega GmbH, Wiesloch, Baden Württemberg, Germany) but instead of boiling proteins for 5 minutes at 95°C we performed the denaturation step at 50°C for 30 minutes to maintain the integrity of the protein. SDS-polyacrylamide gel electrophoresis and western blotting was performed as previously reported (27) with few modifications. The membrane proteins were blotted on a nitrocellulose membrane for 90 minutes at 100 V with an additional ice block at 4°C. To mask unspecific binding sites the membrane was blocked with BlueBlock PF (Serva, Heidelberg, Germany). The split membranes were incubated with the primary monoclonal antibodies α5, 12G10 anti-α-tubulin (DSHB Hybridoma Bank, University of Iowa; deposited by D.M. Fambrough (43) and by J. Frankel and E.M. Nelsen), and customized specific polyclonal antibodies for the subunits β1, β2, and β3 (Davids Biotechnology, Regensburg, Germany). For detection of the primary antibodies the membrane was incubated for one hour with goat anti-mouse (α5, 12G10 anti-α-tubulin), anti-rabbit (β2), and anti-chicken (β1, β3) secondary antibodies conjugated with horseradish peroxidase (Dianova, Hamburg, Germany) and developed with 4-chloro-1-naphtol as previously described (27).

### Extraction and preparation of cardenolides

Calotropin was extracted according to (35) from approximately 8 g of dried larvae of the monarch butterfly *Danaus plexippus* reared on *Asclepias curassavica*. The cardenolides were extracted with methanol, lipids extracted with hexane, and the cardenolides retrieved by chloroform extraction. The concentrated methanolic extract was subjected to preparative HPLC (Agilent 1100) equipped with a photodiode array detector and a semi-preparative RP-Nucleodur C18 Htec column (5 µm, 250 × 10 mm, Macherey-Nagel, Düren, Germany) using a water-acetonitrile gradient. Calotropin was identified by its retention time and checked for molecular weight and LS-MS spectrum by LC-ESI-TOF spectrometry by Dr. Maria Riedner (Department of Chemistry, Universität Hamburg) and the structure confirmed by NMR spectroscopy. The resulting calotropin preparation was 80% pure.

Calotropin and the commercially available ouabain (O3125, Sigma-Aldrich) were first dissolved in DMSO and prepared for Na,K-ATPase activity assays as dilutions in sterile H_2_O reaching final concentrations from 10^−8^ to 10^−3^ M with 2% DMSO.

### Na,K-ATPase activity and inhibition assay

Activities of the nine different Na,K-ATPase α/β-subunit combinations of *O. fasciatus* were determined by photometric measurements of inorganic phosphate released from enzymatic ATP hydrolysis while subtracting background ATPase activity following previously described methods (27,44). Based on previous results (27) the buffer regime used was 50 mM NaCl, 4 mM MgCl_2_, 50 mM imidazol, 20 mM KCl, pH 7.4. The inhibition of the different Na,K-ATPases was tested under increasing ouabain and calotropin concentrations and expressed as percentage activity of a non-inhibited control. For each α/β-subunit construct technical duplicates of three biological replicates of recombinant proteins isolated from independently transfected *Sf*9 cells were tested.

### Statistic analysis of data

After checking for normal distribution (Shapiro Wilk’s test) and variance homogeneity (Levene’s test on squared deviations), differences in total Na,K-ATPase activities were tested and compared between each α/β-subunit construct by one-way ANOVA followed by Tukey’s HSD posthoc test. The data of each Na,K-ATPase α/β-subunit construct was subjected to nonlinear curve fitting with dose response function and Levenberg Marquardt iteration algorithm in Origin-Pro 9.1G (OriginLab Corporation, Northampton, USA) using a four parameter logistic function, top asymptote was set to 100, bottom asymptote set to 0 and a variable Hill slope. For each construct IC_50_ values were calculated and differences were analyzed separately for constructs affected by ouabain and calotropin by one-way ANOVA followed by Tukey’s HSD posthoc tests. A pairwise, F-test based comparison of similarity between the fitted curves was performed using the command “compare datasets”. All calculations were performed with Origin-Pro 9.1G.

## Declarations

### Ethics approval and consent to participate

Not applicable.

### Consent for publication

Not applicable.

### Availability of data and materials

The datasets generated during the current study are available in the Dryad repository at https://doi.org/10.5061/dryad.j0zpc86dg, sequence data will be deposited in the European Nucleotide Archive (ENA) upon acceptance.

### Author’s contribution

SDa and SDo conceptualization; SDa, MW, VW, RT, MH cloning and protein expression; SDa and MW enzyme assays, data analysis and visualization; MW and SDo writing-original draft; SDa writing-review and editing.

## Acknowledgments

We acknowledge the help of Sebastian Gatemann who purified calotropin as part of his BSc thesis. Tammy Tischer supervised by Jennifer Lohr complemented the 5’ end of the αC copy. Shab Mohammadi improved the manuscript by proofreading the English and valuable discussions. We thank Daniel Tal for generating the structures in Fig. 1.

## Supplemental

**Figure S1:**
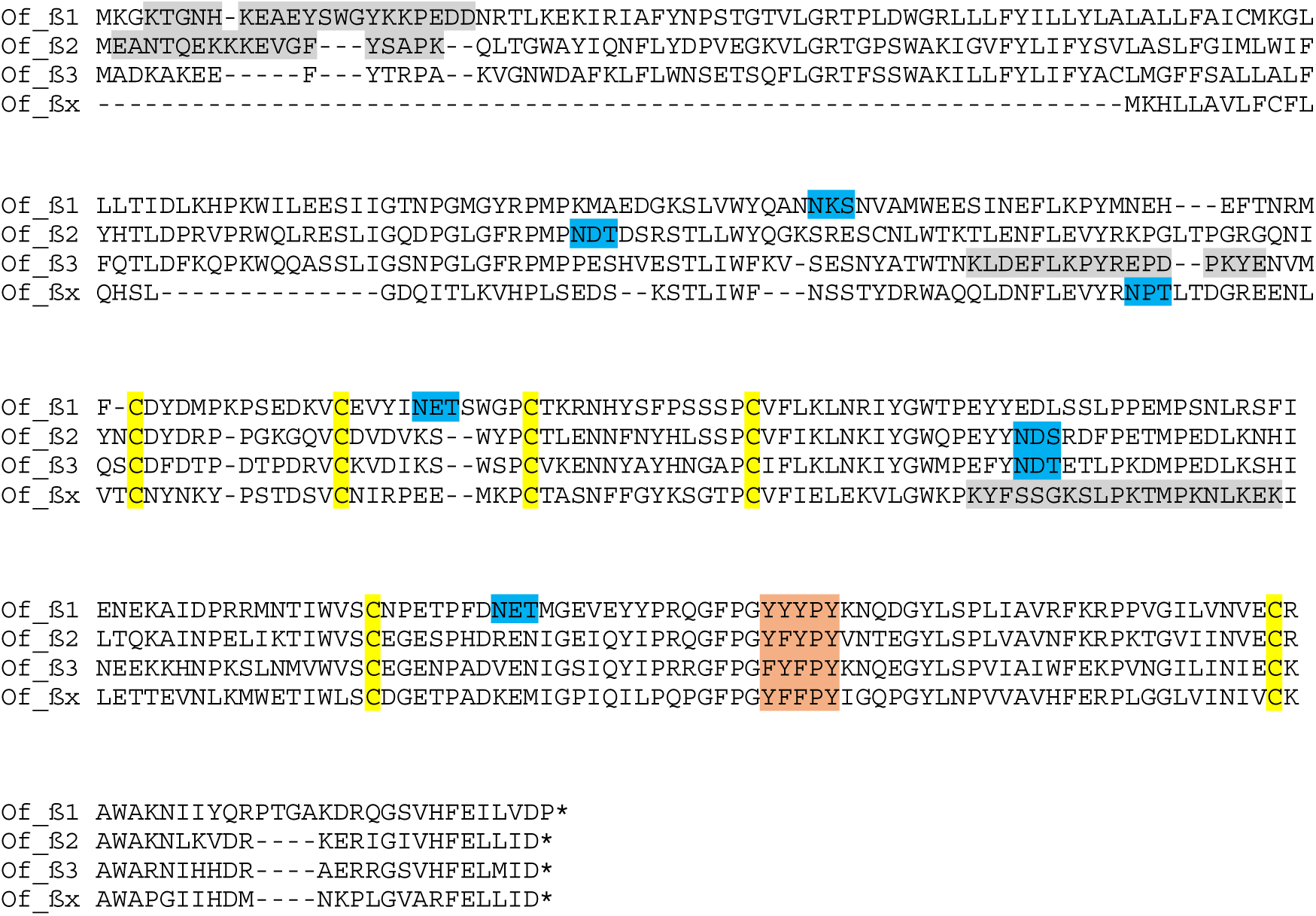
Alignment of deduced *O. fasciatus* Na,K-ATPase α and β-subunit protein sequences. Full length coding sequence of all four β-subunits, tyrosine rich motif typical for β-subunits in orange, cysteine residues forming three conserved intramolecular disulfide bridges in yellow, potential glycosylation sites in blue, and peptides used to raise β-subunit specific antibodies are marked in grey. The N-terminal truncated βx was not further investigated in this study.

**Figure S2:**
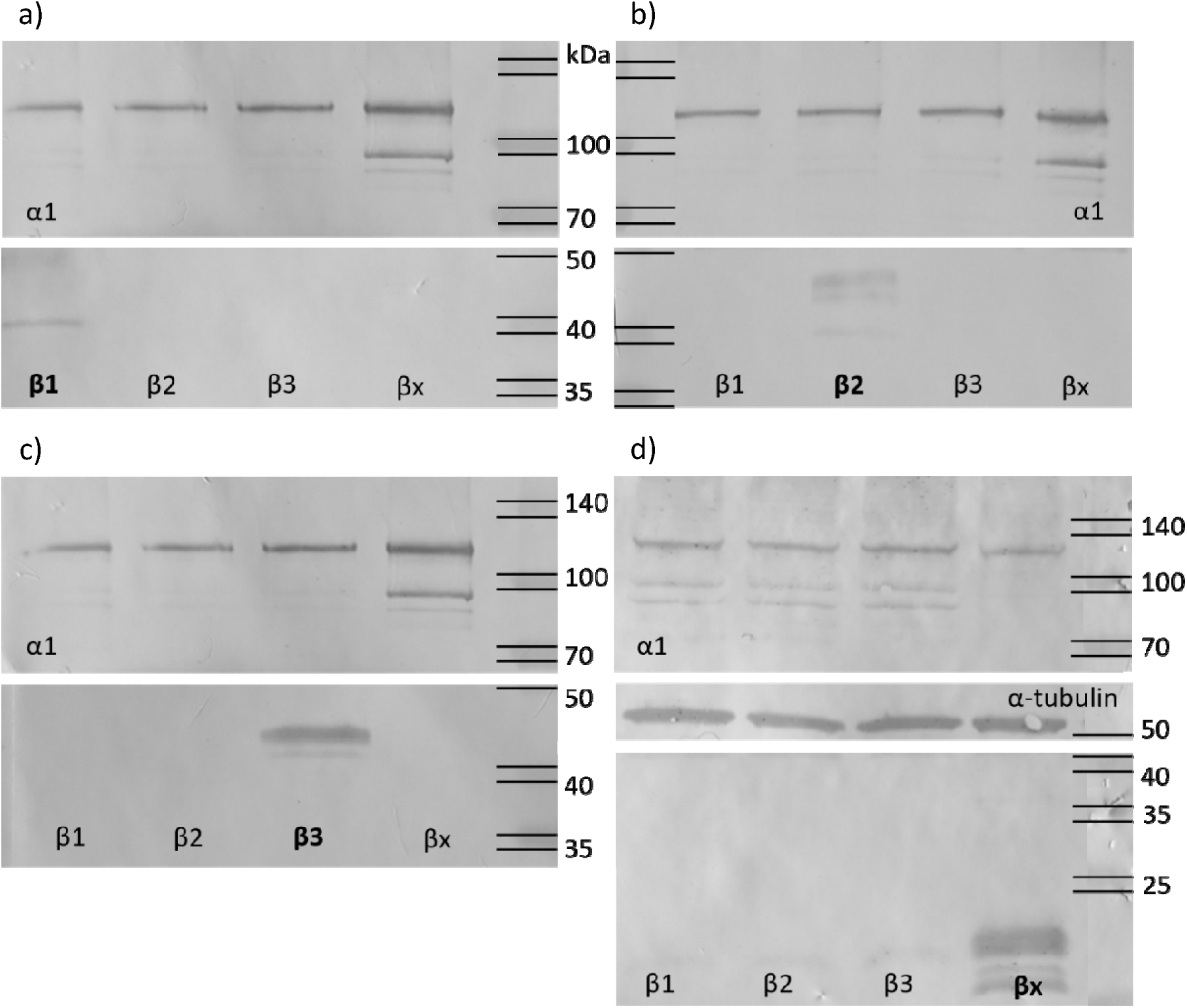
Antibody specificity test with 30µg of heterologously expressed α/β enzymes via SDS-PAGE and western blot. Each set of α1 combined with β1, β2, β3, and βx respectively was stained with α5 primary antibody (upper membrane with α1-subunits at 110 kDa) and one of the four β-specific antibodies: a) β1, b) β2, c) β3, and d) βx-specific antibody, lower membrane with dissociated β-subunits). The middle membrane of the last set (d) was additionally stained with 12G10-anti-α-tubulin. Both, the α-tubulin and α1-staining function as a positive control. The primary antibodies were detected with HRP-conjugated secondary antibody (α5 and 12G10-anti-α-tubulin: goat-anti mouse, β1 and β3: goat-anti chicken, β2 and βx: goat-anti rabbit). The western blots show that the β-specific antibodies solely stain their target subunits (bold letters mark the specific stained β-subunits).

**Figure S3:**
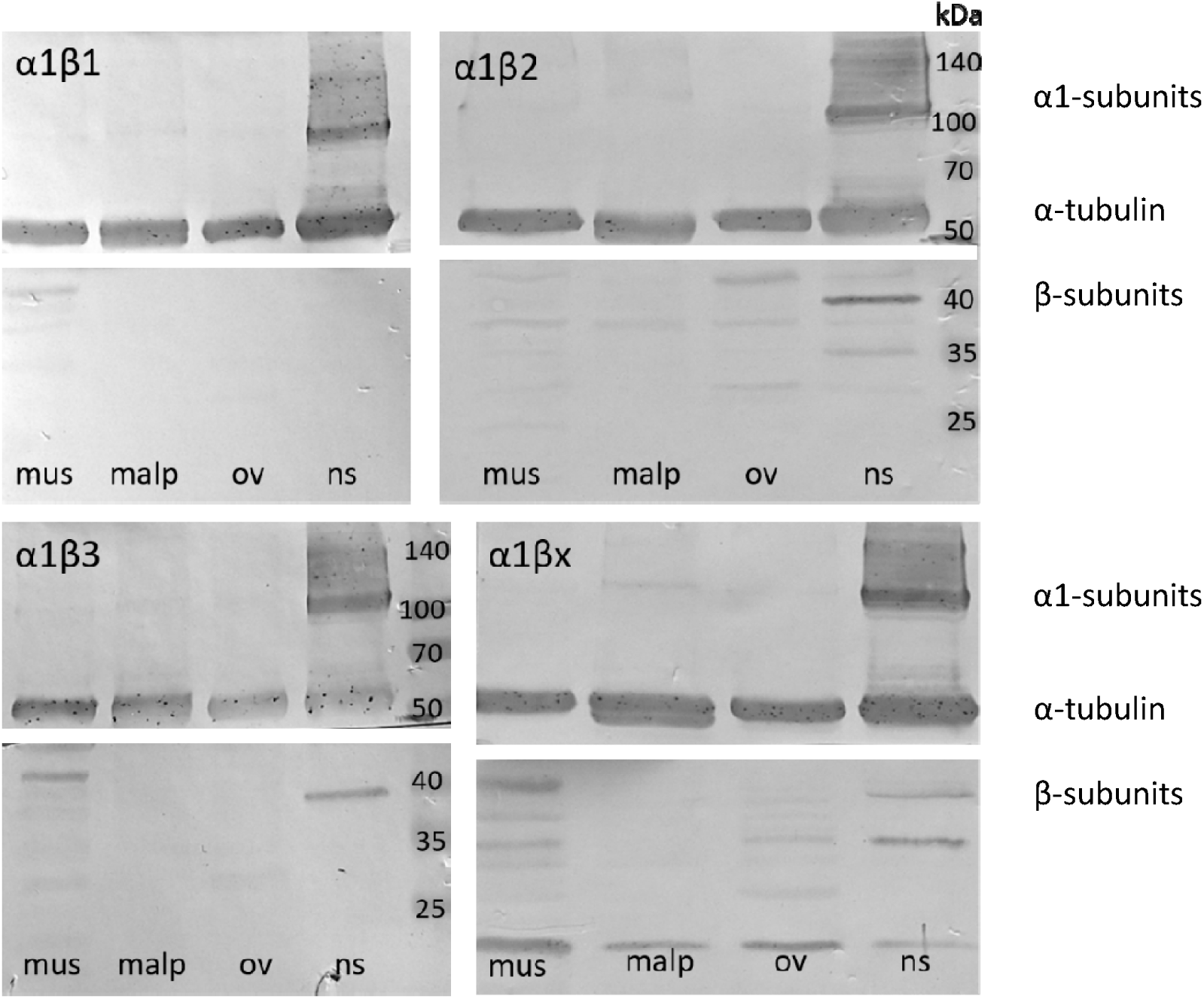
**Antibody specificity test with 35µg four different tissue extracts (muscle (mus), Malpighian tubules (malp), ovaries (ov), nervous tissue (ns)) via SDS-PAGE and western blot.** On each membrane set, consisting of an upper and lower slice, one α1/β-complex is shown. Each set of four different tissue extracts was stained with α5 primary antibody (α1: 110 kDa) and 12G10-anti-α-tubulin (α-tubulin: 55 kDa) on the upper slices and one of the four β-specific antibodies (β: 30-45 kDa; from upper row left to bottom row right: β1-, β2-, β3- and βx-specific antibody) on the lower slices. The 12G10-anti-α-tubulin staining functions as a positive control. The primary antibodies were detected with HRP-conjugated secondary antibody (α5 and 12G10-anti-α-tubulin: goat-anti mouse, β1 and β3: goat-anti chicken, β2 and βx: goat-anti rabbit). The immunostained western blots show differences between abundances and glycosylation pattern in each tissue unique for each β-subunit.

